# Sub-lethal exposure to chlorfenapyr reduces the probability of developing *Plasmodium falciparum* parasites in surviving *Anopheles* mosquitoes

**DOI:** 10.1101/2023.07.03.547458

**Authors:** Prisca A. Kweyamba, Lorenz M. Hofer, Ummi A. Kibondo, Rehema Y. Mwanga, Rajabu M. Sayi, Fatuma Matwewe, James W Austin, Susanne Stutz, Sarah J Moore, Pie Müller, Mgeni M. Tambwe

## Abstract

Pyrethroid resistance in the key malaria vectors threatens the success of pyrethroid-treated nets. To overcome pyrethroid resistance, Interceptor® G2 (IG2), a ‘first-in-class’ dual insecticidal net that combines alpha-cypermethrin with chlorfenapyr was developed. Chlorfenapyr is a pro-insecticide, requiring bio-activation by oxidative metabolism within the insect’s mitochondria, constituting a mode of action preventing cross-resistance to pyrethroids. Recent epidemiological trials conducted in Benin and Tanzania confirm IG2’s public health value in areas with pyrethroid-resistant *Anopheles* mosquitoes. As chlorfenapyr might also interfere with the metabolic mechanism of the *Plasmodium* parasite, we hypothesised that chlorfenapyr may provide additional transmission-reducing effects even if a mosquito survives a sub-lethal dose. Therefore, we tested the effect of chlorfenapyr netting to reduce *Plasmodium falciparum* transmission using a modified WHO tunnel test with a dose yielding sub-lethal effects. Pyrethroid-resistant *Anopheles gambiae s*.*s*. with established mixed-function oxidases and *Vgsc*-L995F knockdown resistance alleles were exposed to untreated netting and netting treated with 200 mg/m^3^ chlorfenapyr for 8 hours overnight and then fed on gametocytemic blood meals from naturally infected individuals. Prevalence and intensity of oocysts and sporozoites were determined on day 8 and day 16 after feeding. Both prevalence and intensity of *P. falciparum* infection in the surviving mosquitoes were substantially reduced in the chlorfenapyr-exposed mosquitoes compared to untreated nets. The odds ratios in the prevalence of oocysts and sporozoites were 0.33 (95% confidence interval; 95% CI: 0.23-0.46) and 0.43 (95% CI: 0.25-0.73), respectively, while only the incidence rate ratio for oocysts was 0.30 (95% CI: 0.22-0.41). We demonstrated that sub-lethal exposure of pyrethroid-resistant mosquitoes to chlorfenapyr substantially reduces the proportion of infected mosquitoes and the intensity of the *P. falciparum* infection. This will likely also contribute to the reduction of malaria in communities beyond the direct killing of mosquitoes.

**Author summary:** Malaria remains a serious problem in many tropical and sub-tropical areas, affecting the welfare and health of many individuals. Since 2016, malaria has increased and the emergence of mosquitoes that are resistant to different classes of insecticides used in vector control tools may have contributed to some of this increase. Therefore, insecticides with a different mode of action are required to manage vector resistance to insecticides used for public health vector control. One of the main resistance mechanisms is metabolic resistance where mosquitoes upregulate detoxification enzymes to break down insecticides. Chlorfenapyr is a pyrrole-pro-insecticide that is metabolised by these detoxification enzymes from chlorfenapyr to tralopyril that disrupts mitochondrial function in mosquitoes. We therefore hypothesized that the metabolites of chlorfenapyr may also have an effect on *Plasmodia* since they, too possess mitochondria and this could reduce the development of *Plasmodium* in mosquitoes that survived a sub-lethal dose of chlorfenapyr. In this study we established and evaluated a modified WHO tunnel assay to investigate the effect of chlorfenapyr in *Plasmodium*-infected *Anopheles* mosquitoes. In this bioassay, we found that chlorfenapyr substantially reduces the proportion of *Plasmodium*-infected mosquitoes at doses sub-lethal to mosquitoes. Our findings demonstrate that chlorfenapyr provides additional benefits beyond mosquito killing although the mechanism of action requires further elucidation.

## Introduction

Combining active ingredients (AI) with two different modes of action on insecticide-treated nets (ITNs) or by using both ITNs and indoor residual spray (IRS) together with different active ingredients is beneficial for the control of insecticide-resistant mosquitoes in endemic settings [1]. One new tool that has recently been recommended for public health is Interceptor® G2 (IG2; BASF SE, Ludwigshafen, Germany) which includes a pyrethroid and chlorfenapyr, a pro-insecticide that is designed to enhance for resistance management [2]. Recent trials of IG2 have demonstrated significant reduction in malaria prevalence compared to pyrethroid only ITNs of 55% in Tanzania [3] and 46% in Benin [4]. There are also multiple operational trials that demonstrate the improved control of malaria using IG2 ITNs in comparison to pyrethroid only ITNs [5]. Pyrethroid-chlorfenapyr ITNs have recently been recommended for use in the control of malaria areas of malaria transmission where mosquitoes are resistant to pyrethroids [1].

Chlorfenapyr is an Insecticide Resistance Action Committee (IRAC) Group 13 insecticide: a pyrrole chemistry. It uncouples oxidative phosphorylation via disruption of the proton gradient as a protonophore to short-circuit mitochondrial respiration through inner-mitochondrial membranes of insect cells so that adenosine triphosphate (ATP) cannot be synthesised and subsequently robs insects of energy resulting in death [2, 6]. While parent chlorfenapyr (CL303630) can be relatively low in toxicity to mosquitoes for some testing modalities, chlorfenapyr requires metabolisation to its pro-insecticidal metabolite, tralopyril (CL303268), in the mosquitoes to elicit lethal effects [6]. This process is dependent on mosquito metabolism, a process that may take time to begin, but once conversion is started the insect’s respiration is increased which then enhances additional conversion of the parent chlorfenapyr to tralopyril [6]. In nature, mosquitoes will encounter chlorfenapyr while metabolically active, foraging at night when chlorfenapyr is applied to ITNs or after being physiologically or metabolically active from foraging, or while resting on treated walls for blood digestion if clorfenapyr is applied as an IRS. It is also likely that further conversion to the metabolite will occur while gravid mosquitoes are again metabolically active while seeking a place to oviposit.

Metabolic resistance is one of the main mechanisms of pyrethroid-resistance observed in malaria vectors [7]. As pyrethroids are used almost ubiquitously for malaria control, metabolic pyrethroid resistance is widespread throughout sub-Saharan Africa and to a lesser degree to other malaria endemic areas [8]. Metabolic resistance is where the production enzymes of one or several detoxification gene families, such as cytochrome P450-dependent monooxygenases (P450s), carboxyl-cholinesterases (COEs), and/or glutathione S-transferases (GSTs) are increased and used by mosquitoes to sequester, break down or export insecticides so they can survive exposure to lethal concentrations [9]. While this metabolism is a detoxification process, it can increase the potency of a pro-insecticide that is metabolised from a parent molecule and may, therefore, be exploited as a means to control metabolically resistant insect populations [2].

It is known that exposure to doses of insecticides that are sub-lethal for insecticide-resistant mosquitoes can interfere with parasite development inside mosquitoes [10], while the underlying mechanisms remain unknown. Owing to the mode of action of chlorfenapyr on mitochondrial respiration in many organisms [6], we hypothesised that chlorfenapyr may also interfere with the metabolic mechanism of the *Plasmodium* parasite in infected *Anopheles* mosquitoes and thereby interfere with malaria parasite development in mosquitoes that were exposed to sub-lethal doses of chlorfenapyr.

For this reason, we developed and evaluated a modified WHO tunnel test allowing to expose mosquitoes to a sub-lethal dose in order to measure whether pre-exposure to chlorfenapyr impacts the prevalence and intensity of infection of *P. falciparum* among *An. gambiae s*.*s*. mosquitoes that survived exposure to chlorfenapyr-treated nets.

## Results

### Selection of chlorfenapyr dose

While the mortality rates were significantly higher in the chlorfenapyr-exposed mosquitoes, the mortality rates between the 100 and 200 mg/m^2^ chlorfenapyr net was similar and below 50% (Table 1). Therefore, the following experiments with infected blood were conducted using the 200 mg/m^2^ chlorfenapyr net samples.

**Table 1:**
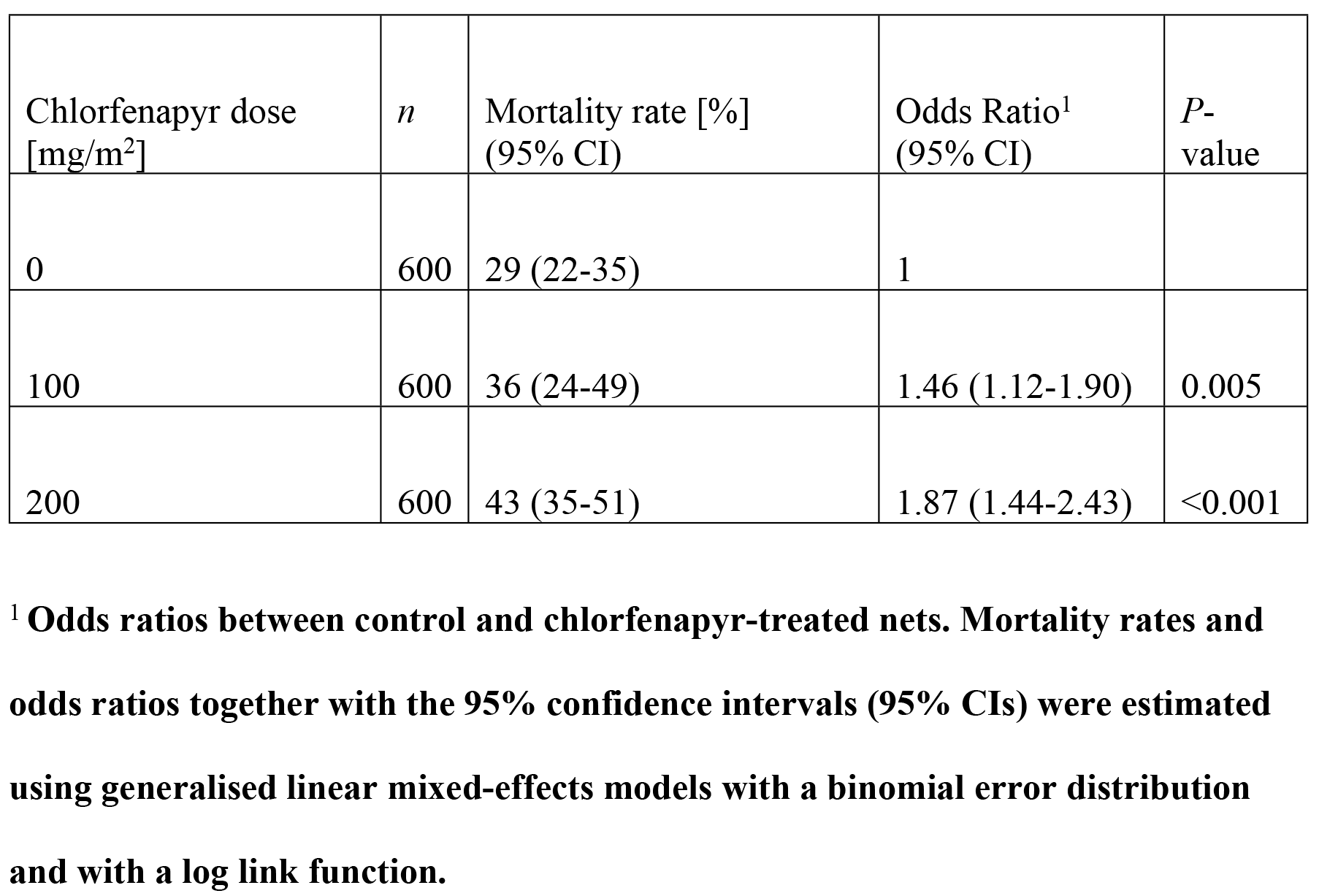
Mortality rates in blood-fed pyrethroid-resistant *Anopheles gambiae s.s*. upon exposure to chlorfenapyr-treated netting.

| Chlorfenapyr dose [mg/m<sup>2</sup>] | *n* | Mortality rate [%] (95% CI) | Odds Ratio<sup>1</sup> (95% CI) | *P*-value |
| --- | --- | --- | --- | --- |
| 0 | 600 | 29 (22-35) | 1 |  |
| 100 | 600 | 36 (24-49) | 1.46 (1.12-1.90) | 0.005 |
| 200 | 600 | 43 (35-51) | 1.87 (1.44-2.43) | <0.001 |

<sup>1</sup> Odds ratios between control and chlorfenapyr-treated nets. Mortality rates and odds ratios together with the 95% confidence intervals (95% CIs) were estimated using generalised linear mixed-effects models with a binomial error distribution and with a log link function.

### Effect of chlorfenapyr on infection rates

Four out of eight individuals tested positive for gametocytes and had more than three gametocytes per 500 white blood cells. The age of these four donors was 6-40 years.

The blood-fed mosquitoes exposed to the net treated with 200 mg/m^2^ chlorfenapyr had much lower oocyst and sporozoite infection rates than those exposed to the untreated control net (Fig 2A & 2B). The odds ratios in the prevalence of oocysts and sporozoites were 0.33 (95% CI: 0.23-0.46) and 0.43 (95% CI: 0.25-0.73), respectively.

**Fig 1:**
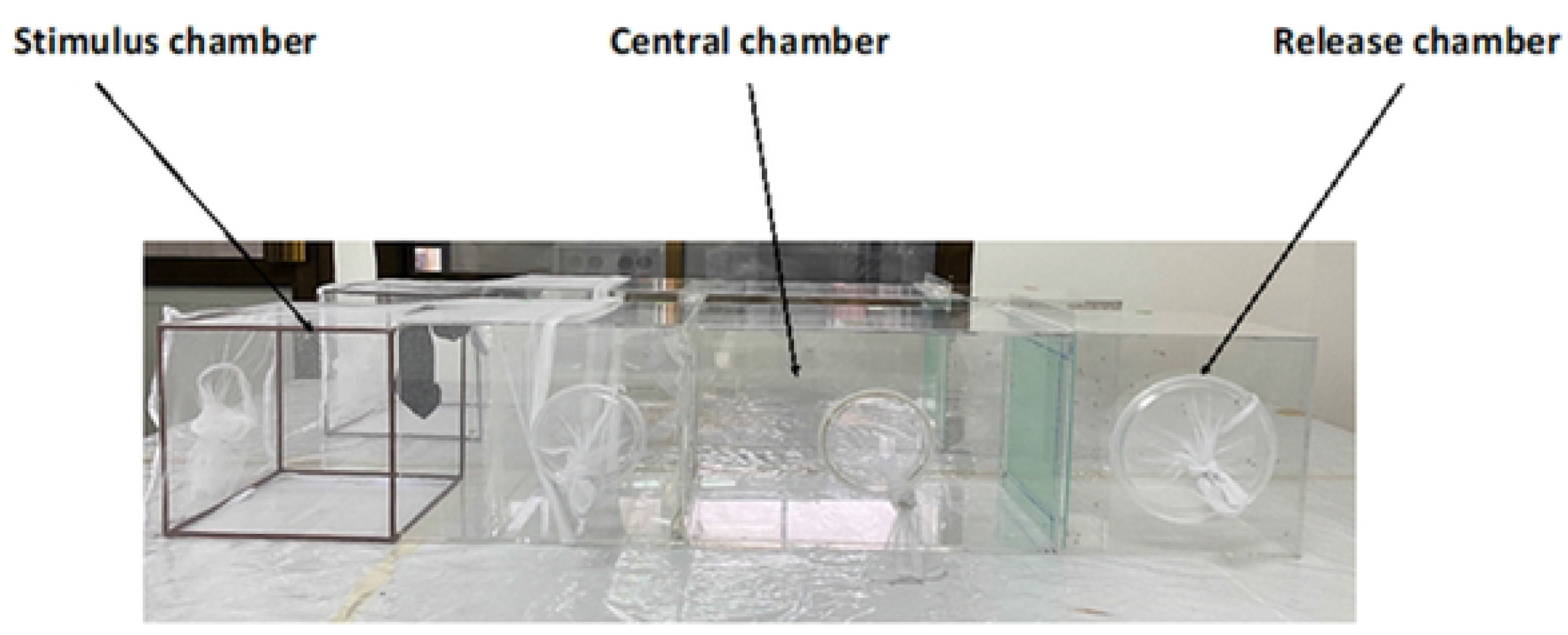
Modified WHO tunnel assay. The main sections are the release, central and stimulus chambers. The partition between the central chamber and the response chamber was designed to attach individual test-treated nets or control. To protect mosquitoes from escaping, the extension cage was made from a Safi net with 100 deniers that do not allow the mosquito to pass and socks were placed in the stimulus chamber as bait to lure mosquitoes towards the test-treated nets or control.

**Fig 2:**
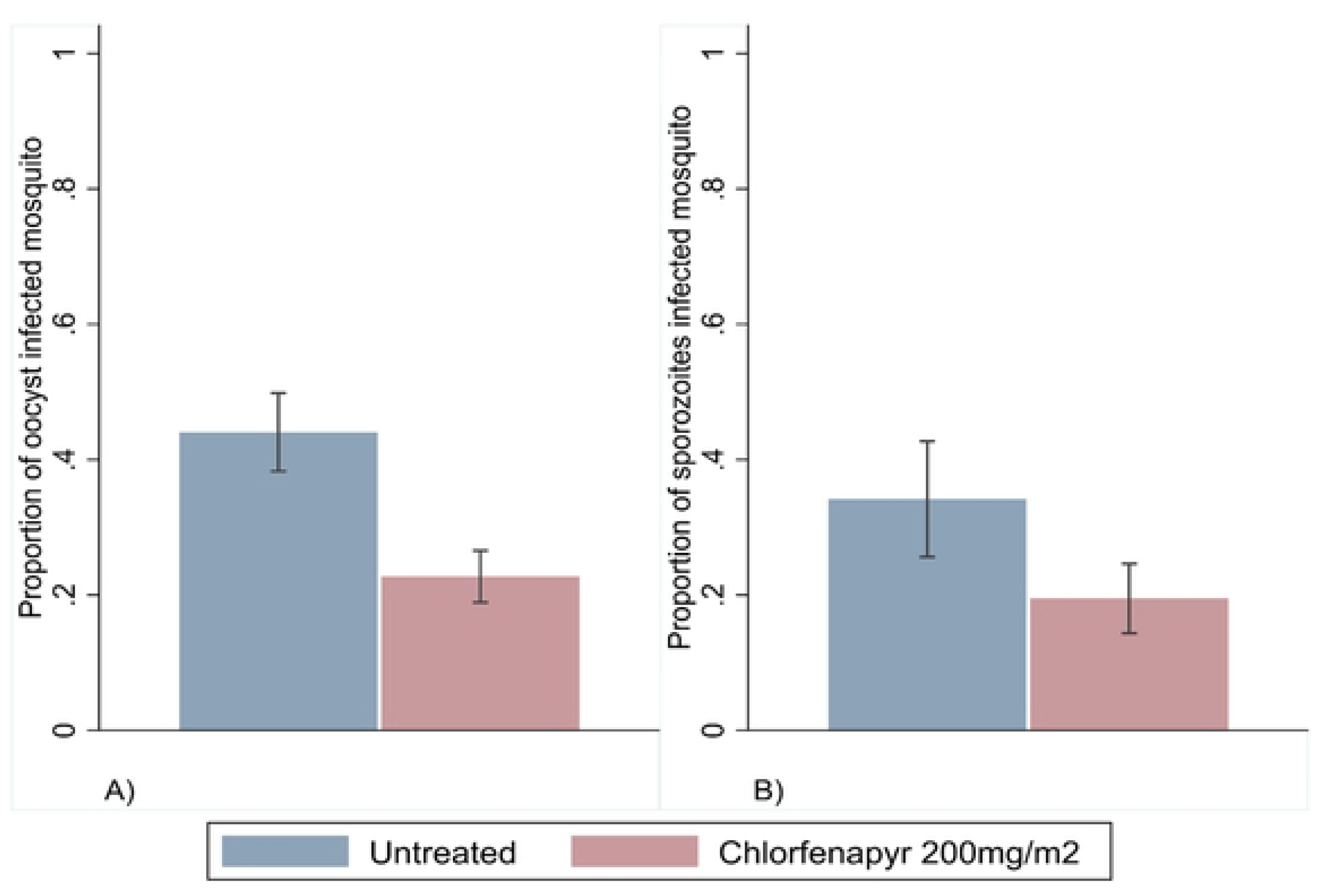
Effects of chlorfenapyr on *Plasmodium falciparum* infection rates in pyrethroid-resistant *Anopheles gambiae s.s*. (A) Oocyst infection rates are determined microscopically. (B) Sporozoite infection rates measured by RT-qPCR.

### Effect of chlorfenapyr on oocyst and sporozoite intensity

In addition to the decreased oocyst infection rate in the chlorfenapyr exposed individuals, the oocyst intensity was also much lower (Fig 3A & 3B). Here, the incidence rate ratio (IRR) was 0.30 (95% CI: 0.22-0.41) and statistically significantly different from 1 (GLMM, *p*-value < 0.003). In contrast, the IRR in the sporozoite rate was statistically not significantly different from 1.

**Fig 3:**
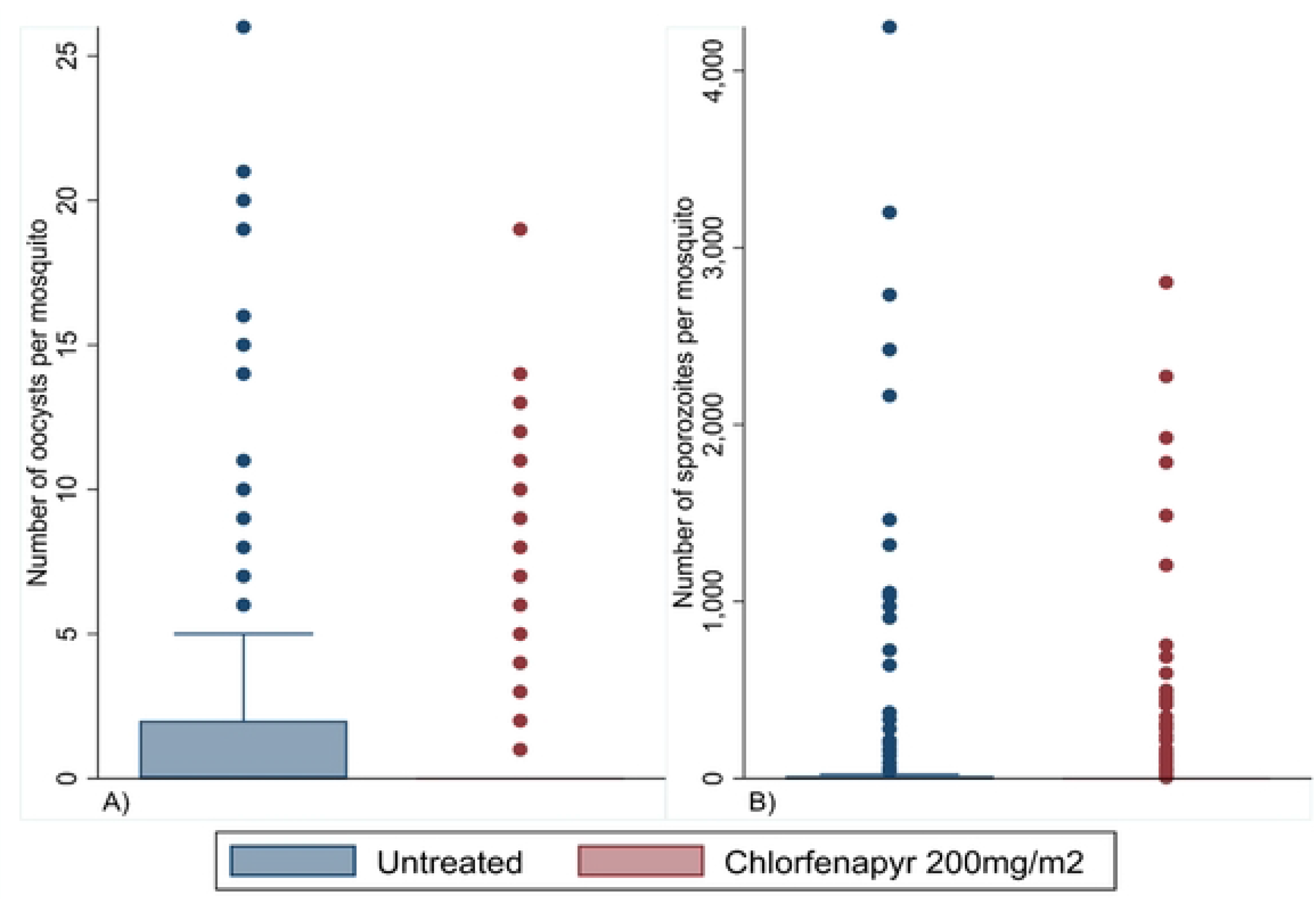
Effects of chlorfenapyr on *Plasmodium falciparum* infection intensities in pyrethroid-resistant *Anopheles gambiae s.s*. (A) Oocyst infection intensity was determined microscopically. (B) Sporozoite infection intensity measured by RT-qPCR.

## Discussion

These results describe for the first time an effect of a chlorfenapyr-treated net on the development of wild malaria parasites in pyrethroid-resistant *Anopheles* mosquitoes. In this study we have used a modified WHO tunnel assay to pre-expose mosquitoes to a chlorfenapyr-treated net and have them actively flying, allowing to study the impact of chlorfenapyr on malaria parasites at sub-lethal exposure. When the exposed mosquitoes were subsequently offered a blood meal containing malaria gametocytes from blood donors in Tanzania, half of the mosquitoes did not develop oocysts or sporozoites, despite the fact that the mosquitoes survived the extrinsic incubation period.

Exposure of mosquitoes to the net treated with 100 mg/m^2^ or 200 mg/m^2^ chlorfenapyr induced similar mortality rates. Therefore, experiments were conducted using 200 mg/m^2^ chlorfenapyr dose. It was advantageous to evaluate 200mg/m^2^ of chlorfenapyr because this reflects the amount present in the IG2 mosquito net that has alpha-cypermethrin (100 mg/m^2^) and chlorfenapyr (200 mg/m^2^) coated onto polyester netting. The results from this experiment reflected likely concentrations that would be encountered under field conditions. However, neither nets with pyrethroid nor a combination between pyrethroid and chlorfenapyr were included as these have also been shown to reduce the prevalence and intensity of infection among mosquitoes exposed to sublethal concentrations [10].

Bioassay design is an important factor when evaluating chlorfenapyr and, therefore, tests with ‘free-flying’ mosquitoes conducted at night are more appropriate for evaluation of chlorfenapyr products as they predict the results of gold standard experimental hut trials better. Indeed, hut trials generally demonstrate high efficacy against metabolically resistant mosquitoes in both East [11-15] and West Africa [16-21] as well as against *Aedes aegypti* in Mexico [22], while cone bioassays or CDC bottle assays, for example, where mosquitoes remain inactive show poor performance of chlorfenapyr[15, 23-25]. The use of a resistant strain is also clearly important since we have observed little difference in mortality between pyrethroid only ITNs and IG2 ITNs [26] when susceptible strains were used, as has also been seen by several other authors [11, 20]. The enzymatic transformation of chlorfenapyr to tralopyril is dependent on mosquito metabolism [6] and in nature, mosquitoes will encounter chlorfenapyr while actively host-seeking at night. The modified WHO tunnel test, therefore, allows mosquitoes to fly around at night when their metabolic enzymes are upregulated [27] ensuring greater conversion of chlorfenapyr to tralopyril.

While several studies have shown a reduction in parasite prevalence or intensity due to sublethal exposure of resistant mosquitoes to bendiocarb, DDT [28] or pyrethroids [10, 29] the mechanism for this is not clear. Possible mechanisms could be that the insecticide modifies mosquito physiology such as increasing oxidative stress leading to increased susceptibility insecticide susceptibility or inhibiting the development of the parasite [28, 30, 31]. However, the principal metabolite of chlorfenapyr through n-dealkylation was known to elicit lethal effects on *Plasmodium in vitro* [32]. Similar evidence has been demonstrated where specific anti-malarial drugs such as atovaquone were assessed by *An. gambiae* tarsal contacts to treated surfaces at low doses to demonstrate rapid and complete transmission blocking to *P. falciparum* [33]. The limitations of these earlier studies have been their focus exclusively on laboratory studies and different substrates rather than actual netting used in the manufacture of current or new ITNs. An additional study has shown that direct injestion of chlorfenapyr in a sugar meal significantly reduces the prevalence and intensity of infection in mosquitoes[34], though exposures to chlorfenapyr on ITNs that afford bioactivation of chlorfenapyr via tarsal contacts to elicit mortality and impairments *Plasmodium* is both logistically favoured and a recommended delivery mechanism as outlined by WHO.

This study showed that chlorfenapyr delivered on an ITN substantially reduces the proportion of *Plasmodium*-infected mosquitoes and the intensity of infection at sub-lethal doses. As insecticide-treated nets age, bio-efficacy is reduced due to insecticide being washed from the ITN surface during normal user washing [35]. Therefore, it may be possible that some mosquitoes escape a toxic dose as the insecticidal content of the ITNs wanes after several years under user conditions [36]. The observed effect of chlorfenapyr in reducing parasite prevalence and intensity is particularly useful as there is emerging evidence that insecticide-resistant vectors are more competent to *Plasmodium* [37] either because they are more likely to live longer than their susceptible counterparts that are killed by insecticide [30] or have lower immunity to parasites [38], even though many insecticide resistance mechanisms impose a fitness cost on mosquitoes in the absence of selection pressure [39]. This study using wild *Plasmodium* isolates is the first to demonstrate that vector control products based on chlorfenapyr may have additional malaria control benefits through impacting the parasite and the vector simultaneously.

The effect of chlorfenapyr on *P. falciparum* observed in the present study could also explain to some extent why IG2 ITNs showed additional efficacy as compared to ITNs that contain a combination of a pyrethroid and PBO or a pyrethroid and pyriproxyfen. Although it is conceivable that chlorfenapyr directly affects *P. falciparum* in mosquitoes exposed to the insecticide, the observed phenomenon may also be due to indirect effects, such as increased insecticide susceptibility in infected mosquitoes, or a combination of direct and indirect mechanisms. Therefore, additional studies are required to further explore the nature of this mechanisms and chlorfenapyr’s overall ability to affect malaria transmission.

In conclusion, we demonstrated that sub-lethal exposure of pyrethroid-resistant mosquitoes to chlorfenapyr substantially reduces the proportion of infected mosquitoes and the intensity of the *P. falciparum* infection. This will likely also contribute to the reduction of malaria in communities beyond the direct killing of mosquitoes.

## Material and methods

### Ethics statement

Written informed consent was obtained from all adult study participants prior to commencement of the study. For children under 18 years their parent or guardian consented and the children assented to participate. All study volunteers were provided with artemisinin lumefantrine treatment within 24 hours of malaria diagnosis as per Tanzania Guidelines for Diagnosis and Treatment of Malaria [40], administered by a medical officer. No adverse effects were reported among the volunteers throughout the duration of the study. Study activities were approved by the Institutional Review Board (IHI IHI/IRB/No: 04-2022) and the National Institute for Medical Research Tanzania (NIMR/HQ/R.8a/Vol.IX/4172).

### Study design

The study was a single (statistician) blinded full factoral design using a modified WHO tunnel test to (1) determine the dose of chlorfenapyr on an ITN that was sufficiently low for at least 50% of *An. gambiae s.s*. mosquitoes to survive for up to 9 days, corresponding to the extrinsic incubation period for oocysts, after exposure and (2) measure whether pre-exposure to the chlorfenapyr ITN impacted the prevalence or intensity of oocysts and sporozoites in *An. gambiae s.s*. mosquitoes fed on blood containing *P. falciparum* gametocytes.

### Mosquito rearing

*Anopheles gambiae* s.s. from the Kisumu KDR strain were obtained from the Centers for Disease Control in Kisumu, Kenya in 2018 and maintained in the colony at the Ifakara Health Institute (IHI), Bagamoyo. The strain is partially pyrethroid-resistant with 50% survival of 1x WHO discriminating concentration of permethrin and expression of both voltage-gated sodium channel *Vgsc*-L995F knockdown resistance (*kdr*) and *GST*e*1* mutation (J. Gimnig pers. comm.). This strain was used because previous experiments had demonstrated the strain to have long survival and good development of parasites (Hofer unpublished). Larvae were maintained at a density of 200 per litre of water and fed 0.3 g per larva on Tetramin fish food (Tetra Ltd., UK). For colony maintenance, the adult mosquitoes were fed on cow blood between 3 and 6 days after emergence for egg development using a Hemotek^®^ membrane feeder (SP-6 System, Hemotek Ltd., Blackburn, UK). Mosquitoes were provided with autoclaved 10% sucrose solution *ad libitum*. Temperature and relative humidity within the insectary are maintained between 27±2ºC and 60%-85% following the MR4 guidance [41]. Strict hygiene measures were maintained to ensure the colony was free of microsporidia.

### Experimental nets

Untreated 100 denier multi-filament polyester nets and treated multi-filament polyester nets coated with 100 mg/m^2^ and 200 mg/m^2^ chlorfenapyr were used in the study. Nets were provided by BASF SE (Ludwigshafen, Germany).

### Modified WHO tunnel test

The modified tunnel test set-up was first used in testing the behaviour of mosquitoes around vector control tools by Thiévent *et al*. [42] and was further modified to fit the purpose of the present study. Here, the tunnel measured 25 cm × 25 cm × 122 cm and was made of glass. The tunnel is divided into three sections, including a release chamber, a central chamber and a stimulus chamber (Figure 1). The specific modifications were; (1) the use of worn socks as a stimulus, (2) adding an extension cage (25 cm x 25 cm x 25 cm) with the worn sock inside, (3) removing the partition between the release and central chamber to increase the number of mosquitoes accessing the stimulus chamber.

### Experimental procedures

In preliminary tests, mosquito survival was assessed after exposure to the untreated and chlorfenapyr-treated net samples. Survival was assessed over 9 days after exposure to the net samples. Nine days were chosen because sufficient numbers of chlorfenapyr exposed mosquitoes have to survive long enough for malaria parasites to develop into oocycts, a process that takes around 8 days after feeding. Six replicates were conducted for each dose of chlorfenapyr and a negative control to monitor assay quality. In each replicate, 100 mosquitoes were released from paper cups into the release chamber at 22:00 hrs. The following morning at 06:00 hrs, mosquitoes from all chambers were collected and aspirated into paper cups in batches of 50 using a mouth aspirator (J.W. Hock & Co., Florida, USA).

Following the exposure in the tunnel test, mosquitoes were transferred to the feeding room and covered with a black cloth for one hour before receiving a non-infectious blood meal donated by PAK. The non-infectious blood was administered to mosquitoes through water-jacketed glass feeders (14 mm Ø, Chemglass, New Jersey, USA) covered with Parafilm®, connected to a circulating water bath (39°C, ELMI, Newbury Park, CA, USA) via plastic tubing for 15 minutes.

After feeding, the cups were transferred to 30 cm x 30 cm x 30 cm Bugdorm plastic cages (Megaview Science Co., Ltd, Taiwan) and placed in a climatic chamber (S600PLH, AraLab, Lisbon, Portugal). Mosquitoes were left for 48 hours without sugar for unfed mosquitoes to die. Thereafter, mosquitoes were supplied daily with autoclaved 10% sucrose solution on a cotton pad and kept at 75±2% relative humidity and 27±1°C at a 12:12 hours dark:light cycle. Mortality was recorded daily for a over nine days.

Once the optimal experimental netting was identified, the assays were repeated using blood drawn from gametocytemic carriers. Five mililitre of infectious blood was collected from microscopically confirmed gametocytaemic carriers with gametocyte densities greater than three gametocytes per 500 red blood cells. Autologous serum was removed from the whole blood and replaced with pre-warmed malaria-naïve AB serum. Before feeding and after exposure in the tunnel test, mosquitoes were kept in paper cups with access to deionised water for one hour. Then, mosquitoes were allowed to feed for 15 minutes on water-jacketed glass feeders as above. On average, 600 mosquitoes received a blood meal from each participant. Dead mosquitoes were removed from the cup by aspiration 48 hours post blood meal. The mosquitoes were provided with cotton soaked with 10% sucrose solution that was changed daily. Eight days post infection (dpi), one third of the mosquitoes were dissected and midguts were stained using 1% mercurochrome solution before examination for presence of oocysts. The remaining mosquitoes were kept up to 16 dpi and were extracted for molecular diagnosis and quantification of *P. falciparum* infection mosquito stages from the mosquito heads and thoraces (sporozoite stage) using quantitative reverse transcription polymerase chain reaction (RT-qPCR) [43]. While the oocysts were actually counted microscopically, numbers of sporozoites were estimated using a plasmid containing a fragment of the 18S rDNA gene (GenBank: AF145334) from *P. falciparum* (BEI Resources, NIAID, MRA-177). Plasmid copy numbers per μL were calculated as described elsewhere [44] and plasmid standard curves were prepared using serial dilutions over eight magnitudes assuming an average of six copies of the 18s rDNA gene sequence per parasite genome [45]. Each concentration from the serial dilution (standards) was run in triplicate to determine qPCR efficiency, limit of detection, the slope and y-intercept for absolute quantification of *Plasmodium* DNA in sporozoite infected mosquito samples.

### Study participants

The study was conducted in Bagamoyo district located in the coastal region of Tanzania, between September 2021 and August 2022. The target populations were males and females aged 6-40 years from the villages of Wami-Mkoko and Ludiga. Participants who met the inclusion criteria (i.e. asymptomatic, consented 6-40 years old with microscopically detectable gametocytes) were recruited for blood-drawing at the IHI malaria transmission facilities in Bagamoyo. Gametocytes were counted against 500 white blood cells in thick smears. The density of gametocytes were calculated from an estimated leukocyte density of 8,000 per μl of blood.

### Statistical Analysis

Data cleaning and analysis was done in STATA 17 software (StataCorp LLC, College Station TX, USA). Descriptive statistics were used for data summaries, whereby mean mortality and proportion of infected mosquitoes with 95% confidence intervals are presented. For parasite intensity (oocysts or sporozoites), median with minimum and maximum values are presented.

To assess the effect of the low and high dose chlorfenapyr-treated netting on mosquito mortality, mixed-effects logistic regression with a binomial error distribution and a log link function was used. Replicate and dose were included as fixed effects in the model and experimental day was fitted as a random effect.

To assess the effect of sub-lethal chlorfenapyr exposure on *P. falciparum*, mixed-effects logistic regression was used with treatment as a fixed effect and study participants included as a random effect. Prevalence of infected mosquitoes at oocyst and sporozoite stages was evaluated using a generalised linear mixed-effects model with a binomial error distribution and a log link function. For oocyst and sporozoite intensities, models with a negative binomial error distribution and a log link function were fitted.

## Acknowledgements

The authors express their sincere thanks and appreciation to the village leaders and community which surrounds the IHI in Bagamoyo for allowing us to continually run our experiments with minimal interruptions. A special thanks to the Vector Control and Product testing Unit (VCPTU) management, administrators and colleagues who helped in organising logistics and materials, allowing smooth performance of the study. Thanks to Nicolas Brancucci, Matthias Rottmann and Tobias Suter from Swiss TPH for discussions on manuscript contents.

## Notes

### Competing Interest Statement

PAK, SJM, MMT LMH, RMS, RYM, FM and PM conduct product evaluations for a number of companies including BASF. JAW and SS are employed by the BASF Corp, which manufactured chlorfenapyr treated net. All authors have declared that they have no competing interest.

